# Computational approach for decoding Malaria Drug Targets from Single-Cell Transcriptomics and finding potential drug molecule

**DOI:** 10.1101/2024.01.22.576718

**Authors:** Soham Choudhuri, Bhaswar Ghosh

## Abstract

Malaria is a deadly disease caused by *Plasmodium* parasites. While potent drugs are available in the market for malaria treatment, over the years, *Plasmodium* parasites have successfully developed resistance against many, if not all, front-line drugs. This poses a serious threat to global malaria eradication efforts, and the continued discovery of new drugs is necessary to tackle this debilitating disease. With the advent of recent unprecedented progress in machine learning techniques, single-cell transcriptomic in *Plasmodium* offers a powerful tool for identifying crucial proteins as a drug target and subsequent computational prediction of potential drugs. In this study, We have implemented a mutual-information-based feature reduction algorithm with a classification algorithm to select important proteins from transcriptomic datasets (sexual and asexual stages) for *Plasmodium falciparum* and then constructed the protein-protein interaction (PPI) networks of the proteins. The analysis of this PPI network revealed key proteins vital for the survival of *Plasmodium falciparum*. Based on the function and identification of a few strong binding sites on a couple of these key proteins, we computationally predicted a set of potential drug molecules using a deep learning-based technique. Lead drug molecules that satisfy ADMET and drug-likeliness properties are finally reported out of the generated drugs. The study offers a general computational pipeline to identify crucial proteins using scRNA-seq data sets and further development of potential new drugs.

## Introduction

Malaria, caused by Plasmodium parasites, remains a formidable global health challenge. The World Malaria Report 2022 states that there were 619 000 estimated malaria deaths (uncertainty range 577 000–754000) and 247 million estimated cases of malaria (uncertainty range 224–276 million) worldwide in 2021^1^. According to WHO’s report, African Region has the highest malaria burden, with an estimated 95% of cases and 96% of deaths; children under the age of five account for 78.9% of all deaths in this region^1^. Over 200 million cases and nearly half a million deaths are reported annually. Malaria mostly dominates in tropical and subtropical regions, where conditions favor the Anopheles mosquitoes that transmit the malaria parasites. Although malaria is found in regions of Asia, Latin America, the Middle East, and Oceania, it is most common in sub-Saharan Africa^2^. Over time, the treatment of malaria has undergone substantial transformations, with the drugs used to tackle the condition adapting to drug resistance. There were numerous drugs such as Quinine (and its derivatives), Chloroquine, Sulfadoxine-Pyrimethamine (SP), Mefloquine, Artemisinin, and Artemisinin-based Combination Therapies (ACTs), etc, available from 1900 but It’s essential to note that the emergence of drug-resistant variants of the malaria parasite, mainly Plasmodium falciparum, has called into question the efficacy of these drugs^3^. The development of parasite strains that are resistant to drugs has further complicated the treatment procedure, necessitating innovative approaches in the search for effective antimalarial drugs. The incorporation of single-cell transcriptome analysis represents a major development in this endeavor^4^. Malaria parasites can evade the host immune system by modulating host cell gene expression, rendering the infection process extremely flexible and dynamic. While traditional bulk transcriptome methods offer insightful information, they frequently obscure the cellular heterogeneity in infected tissues, resulting in the loss of important data. However, single-cell transcriptomics allows the dissection of this complexity through transcriptome profiles for specific host cells. Plasmodium falciparum has a complicated life cycle involving mosquitoes and human hosts^5–7^. Comprehending this complex cycle is crucial for efficient drug development. The life cycle consists of multiple discrete stages, each with its own set of opportunities and obstacles that can be addressed. The plasmodium life cycle has two main phases: asexual phase(schizogony) in the human host and sexual phase (sporogony) in the mosquito.

### 1. Asexual phase(schizogony)

When an infected female anopheles mosquito bites, sporozoites are injected into the bloodstream and then navigate to the liver^8^. The parasite first infects the liver’s hepatocytes, reproducing schizonts asexually. Then the schizonts burst and Merozoites were released into the bloodstream. Then red blood cells are invaded by merozoites and these merozoites change into a ring stage inside the RBCs.The ring stage develops into a trophozoite, consuming hemoglobin. These trophozoites formed new schizonts and gametocytes. New schizonts can infect more red blood cells, and gametocytes enter mosquitoes when they take blood meals.

### 2. Sexual Phase (Sporogony)

A mosquito consumes gametocytes, or the sexual forms of the parasite when it feeds on an infected human. Gametocytes develop into male (microgametocytes) and female (macrogametocytes) gametes in the mosquito’s intestines^9,10^. A mosquito feeds on blood to fertilize itself, fusing its gametes to form a zygote. On the intestinal wall of the mosquito, the zygote matures into an oocyst. Sporozites are released when oocysts burst and go to the salivary glands of mosquitoes. Targeting genes expressed in gametocytes is essential for interrupting the transmission of the parasite. Drugs designed to inhibit gametocyte development can reduce the spread of malaria in mosquito populations. Identifying genes at every stage of the Plasmodium falciparum life cycle is essential for developing malaria drugs. Focusing on particular genes makes it possible to create drugs that disrupt significant biological processes at particular stages of the parasite’s life cycle. This strategy improves the efficacy of treatments and reduces the possibility of resistance building, which is important in the ongoing battle against malaria. Researchers can create drugs with specific modes of action by comprehending the molecular nuances at each level, ultimately leading to more effective and long-lasting malaria control methods. This study aims to explore the promise of single-cell transcriptomic analysis as a powerful tool for identifying crucial proteins as a drug target and finding suitable drug molecules that can inhibit these proteins (figure 1a). Recent advances in single-cell RNA-sequencing (sc-RNA)techniques paved new ways to characterize gene expression changes during the development stages of the plasmodium life cycle. One of the central advantages of employing sc-RNA methods is the scope of exploring cell-to-cell heterogeneity in the population by uncovering hidden variability in gene expression among single cells. In fact, recent studies have elucidated the role of heterogeneity in enabling a small fraction of the plasmodium population inside the human host to remain ready to enter into the mosquito host by transitioning to the gametogenesis stage. Similarly, heterogeneity plays a crucial role in plasmodium stress response inside the RBC. Moreover, some of these data sets offer information about the stage of each cell in the population, which can be efficiently used to construct several supervised machine-learning models. We have implemented a mutual-information-based supervised feature reduction algorithm and a classification algorithm to select important features from datasets for our analysis. Next, we find some crucial proteins that are important for plasmodium survival using protein-protein interaction network. We find the function and strong binding sites of these crucial proteins. Based on these strong binding sites, we generated drug molecules using a deep learning-based technique. We have used ‘targetdiff’, a non-autoregressive generative diffusion model^11^, and found some lead drug molecules using ADMET and drug-likeliness properties. Next, we did molecular-docking to find the binding energy of selected proteins with their selected ligands. This whole approach allows for identifying subtle variations in gene expression within individual cells, which is crucial for capturing the dynamic responses of different parasite stages. Network-driven analyses in single-cell studies unveil the crucial proteins that are important for plasmodium survival in sexual and asexual stages.

**Figure 1.**
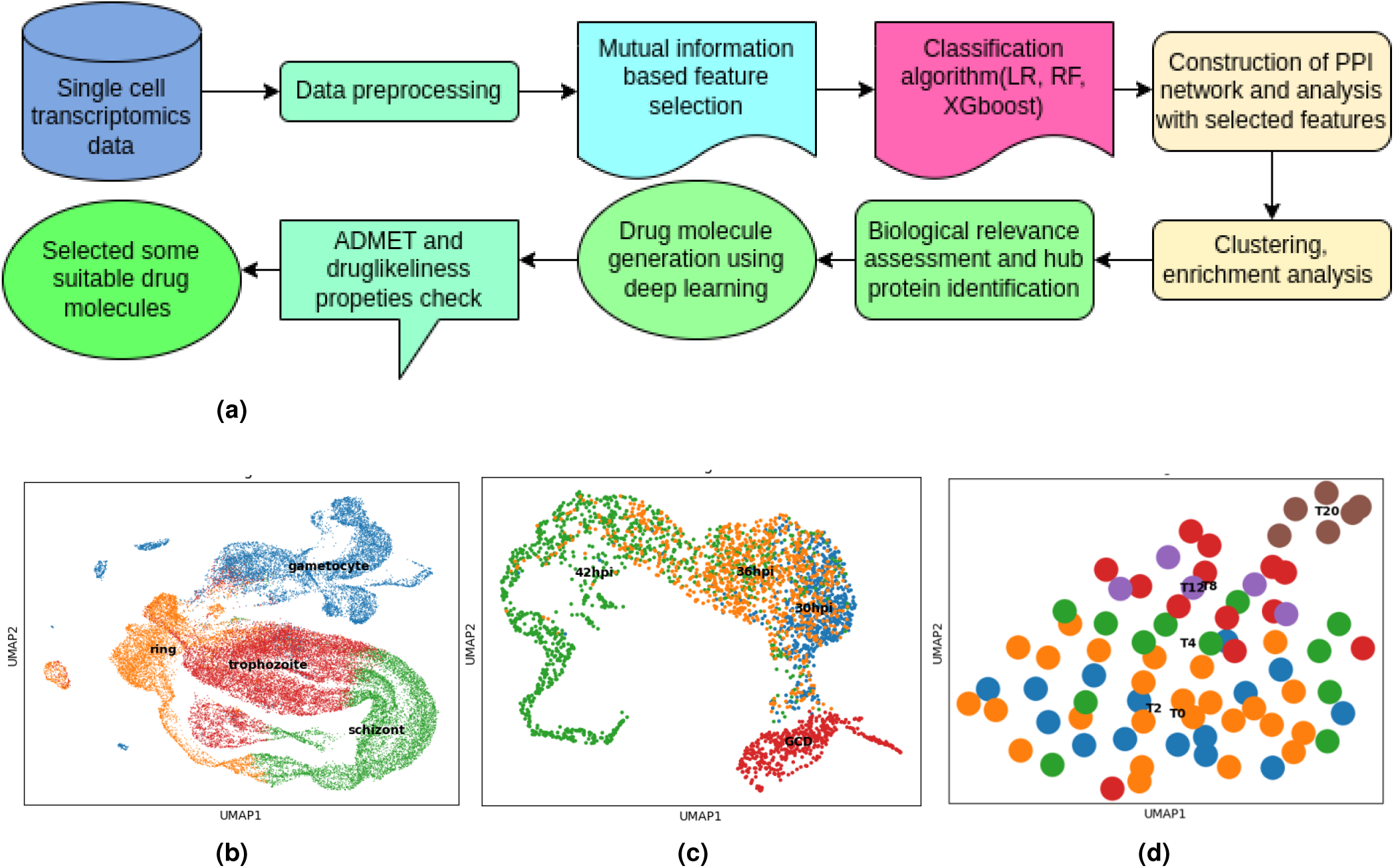
scRNA-seq datasets display distinct expression patterns for cells belonging to different stages. (a) The workflow, describes the steps to select significant features and generate drug molecules. The workflow starts with analysing sc-RNA seq data and selection of features crucial for the developmental stages of *P. falciparum*. Finally deep learning algorithms are utilized to carry out the drug discovery procedure to predict drugs specific to the selected targets. (a) The first dataset contains the expression of 5838 genes for 38635 cells. (b)The Second dataset contains mRNA count of 4722 genes for 3922 cells. (c) Third dataset has 1024 cells and 1024 mRNAs. In first dataset, each cell is assigned one label among the four blood cycle stages (ring, trophozoite, gametocyte, and schizont)^12^. In second dataset, each cell is assigned one label among the four blood cycle stages (30hpi, 36hpi, 42hpi, and GCD)^13^. The dimensional reduction technique UMAP is used to visualize the scRNA-seq data for the (a) 1st (b) 2nd, and (c) 3rd datasets. The cells belonging to different stages are shown with different colors as indicated

## Results

The aim of the study is to identify crucial proteins driving the *Palsmodium falciparum* life cycle stages and finally predicate potential drug molecules which would be able to target the key proteins. The overview of the protocol is described in Figure 1. The protocol consists of several steps ultimately leading to the identification of protein targets and computational prediction of drugs. In the first step, a set of RNA-seq data for both the sexual and asexual phase *Palsmodium falciparum* life cycle is analysed and machine learning and feature selection algorithms were deployed to extract the protein features. Protein-protein interaction network of the features is constructed to finally select crucial proteins out of all the identified features based on network properties which will be discussed in detail in the following sections. In the final step, a deep learning target-based method is employed to predict specific drugs for the selected protein targets.

### The scRNA-seq data clustering reveals distinct developmental stages

We have utilized three publicly available single-cell RNA-seq datasets. Two datasets have been collected from the asexual reproduction phase^12,13^ and one from the sexual reproduction phase^14^. After collecting raw datasets, we preprocessed these datasets, the details of which have been given in the method. The first dataset contains the expression of 5838 genes for 38635 cells. The Second dataset contains mRNA count of 4722 genes for 3922 cells. Third dataset has 1024 cells and 1024 mRNAs. In first dataset, each cell is assigned one label among the four blood cycle stages (ring, trophozoite, gametocyte, and schizont)^12^. In second dataset, each cell is assigned one label among the four blood cycle stages (30hpi, 36hpi, 42hpi, and GCD)^13^. The time points in hours post-invasion (hpi) during the malaria parasite Plasmodium falciparum’s commitment cycle are denoted by the numbers 30, 36, and 42. GCD is the gametocyte stage. The synchronised parasites were cultured, and samples were collected at these precise intervals to examine the transcriptional modifications and alterations in gene expression induced by AP2-G during the commitment cycle^13^. Samples are taken 30, 36, and 42 hours after the invasion, and the time points are measured starting from the beginning of the invasion cycle^13^. In the third dataset, each cell is assigned one label among the four blood cycle stages (T0, T2, T4, T8, T12, T20)^14^. T0, T2, T4, T8, T12 and T20 are time points that represent intervals at which mosquito midguts were collected and dissected for analysis. After taking a blood meal, female Anopheles mosquitoes are tracked in time intervals to check the progression of the infection’s growth. The midguts are taken at the time intervals of 2, 4, 8, 12, and 20 hours after infection to analyze and track the development of the gametocytes or parasites within the mosquitoes^14^. The mRNA counts were RPKM normalized (reads per kilobase of exon per million reads mapped). We have used uniform manifold approximation and projection (UMAP) to get three-dimensional visualization of the cells from the scRNA datasets showing a distinct cluster of different life cycle stages^15^. The distinct life stages of Plasmodium Falciparum are visible before feature selection in Figure 1b, 1c, 1d.

### ML classification models exhibit high accuracy in predicting the developmental stages

Utilization of classification Machine learning algorithm allowed us to predict the blood cycle stage of the parasite cell based on the gene expression pattern. In the initial phase of our study, as described in the previous section, we preprocessed three single-cell transcriptomics datasets, in which every single cell represents one particular developmental stage of the *Plasmodium falciparum*. We first performed data cleaning, normalization, and transformation procedures to ensure a biologically meaningful representation of the datasets. Normalization solves issues such as variability in sequencing depth. A set of classification models(Logistic regression, Random forest, XGboost) for these datasets was implemented to obtain accuracy in predicting the stages. To compute the accuracy, the data sets have been divided into training and test sets following a ratio of 0.8 and 0.2. (Figure 2a). We observed that for all the data sets, the accuracy in all three methods displays high accuracy for both the sexual stages (Figure 3a, 3b blue bars), whereas for the sexual stages, the logistic regression shows higher accuracy compared to the other cases (Figure 3c). This may occur due to this data set’s low number of cells.

**Figure 2.**
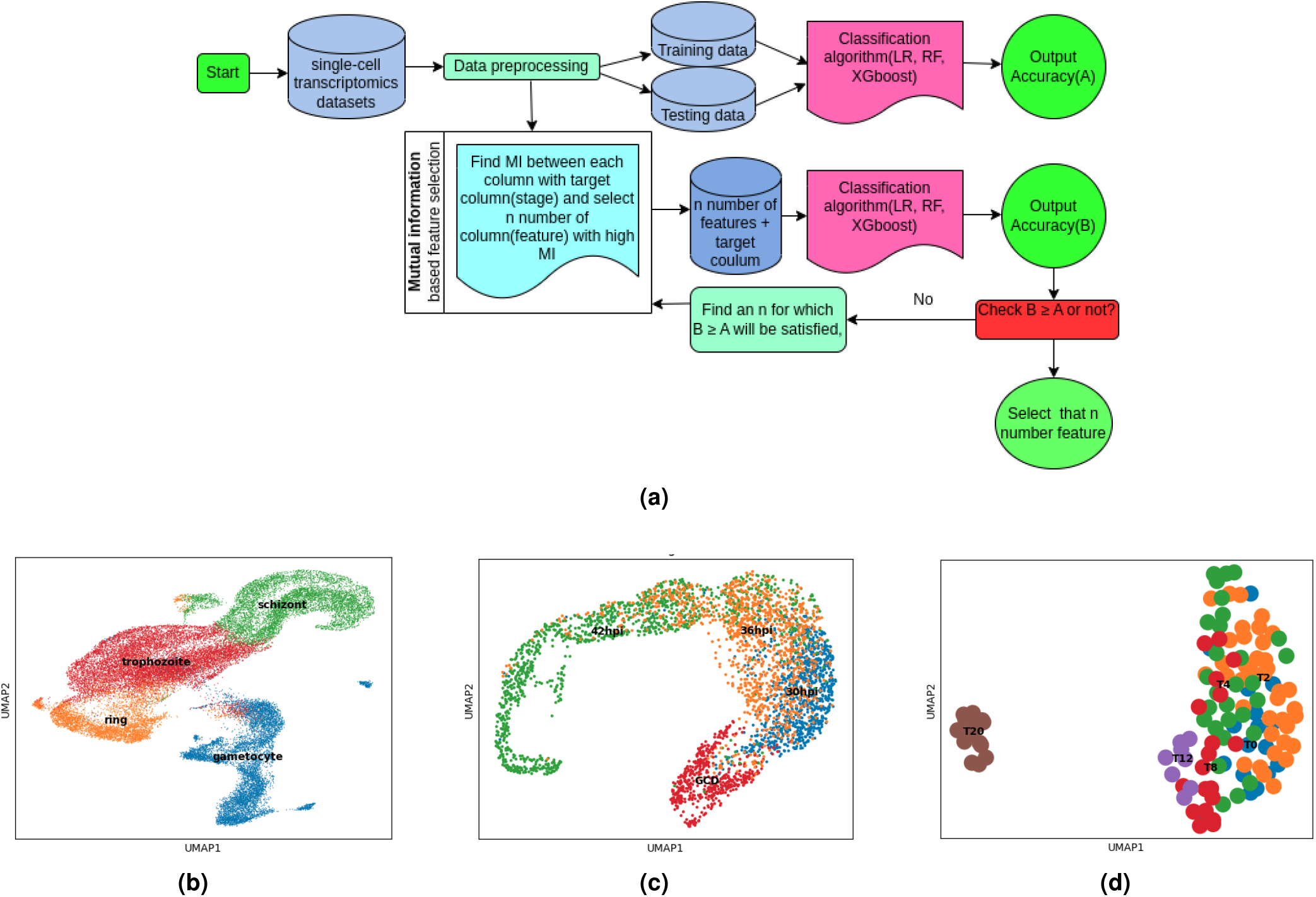
Feature selection algorithm selects key proteins for the developmental cycle. (a) Flowchart of the mutual information-based feature selection pipeline. (b) UMAP of 1st (c) 2nd and (d) 3rd datasets after feature selection

**Figure 3.**
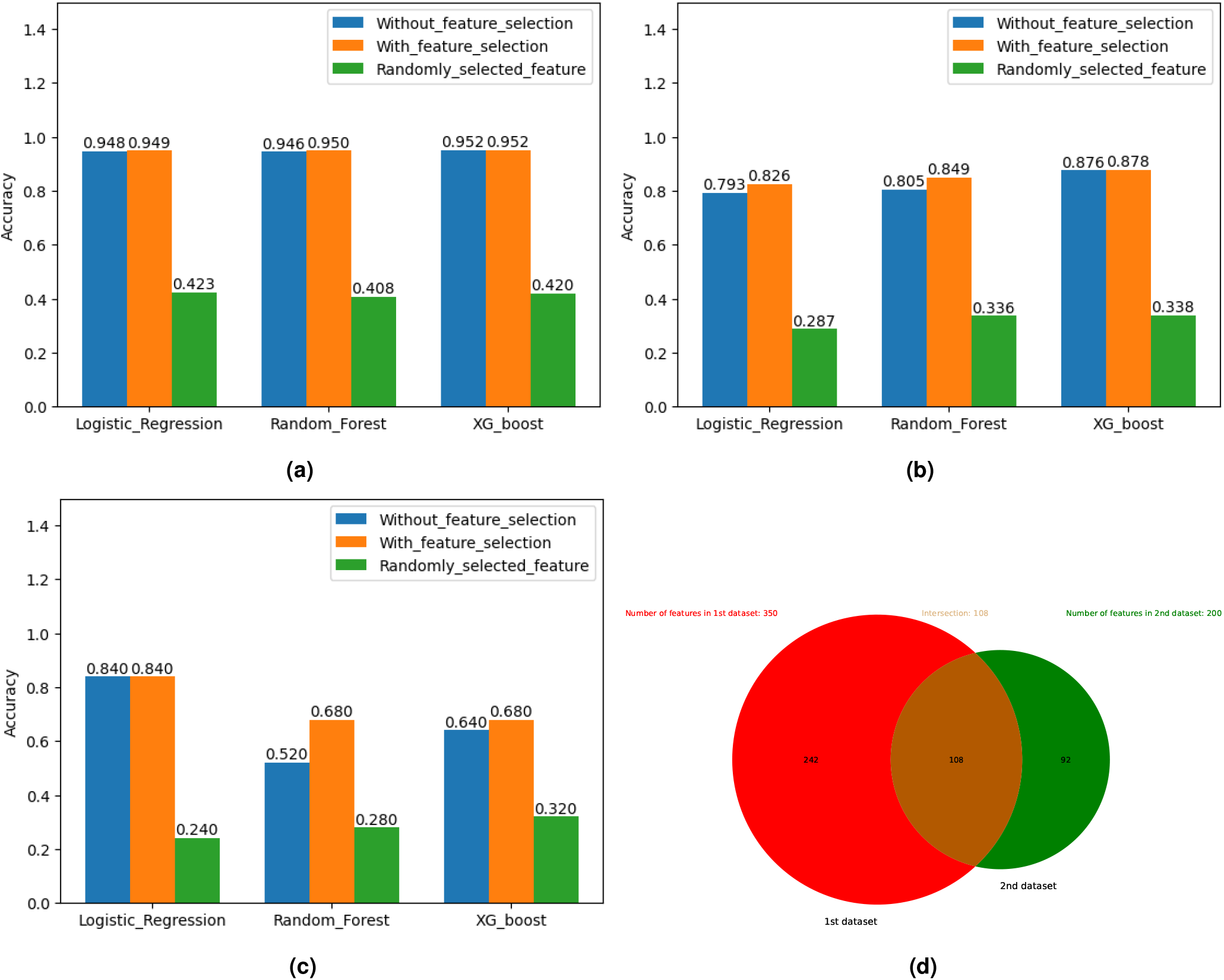
The accuracy of predicting the stages from the gene expression remain almost the same after the feature selection. (a) The accuracy for the first data set before after feature selection and the corresponding accuracy when the features are selected randomly are indicated (b)The accuracy for the second data set before after feature selection and the corresponding accuracy when the features are selected randomly are indicated (c) The accuracy for the third data set before after feature selection and the corresponding accuracy when the features are selected randomly are indicated. (d) The Venn diagram exhibits common selected features among the first and second data sets both for the intra-erythrocyte developmental cycle

### Feature selection algorithm selects key proteins

We implemented a mutual information-based feature reduction technique to select important features from these data sets. The pipeline of this mutual information-based feature reduction algorithm is given in figure 2a. This algorithm calculates mutual information between each column and the target column. The main target of this feature reduction technique is to choose features in such a way that after using these classification models, the accuracy would remain the same or increase. The accuracy of prediction almost remains the same or increases in some cases in both the asexual (Figure 3a, 3b orange bar and sexual stages Figure 3c, orange bar). If the same number of features was randomly selected from these data-sets and applied classification algorithm, then accuracy significantly decreases 3, which implies that these selected features are crucial and contribute more in predicting the stages. The distinct life stages of Plasmodium Falciparum are visible after feature selection in Figure 2b, 2c, 2d. Since data sets for both the two asexual stages are conducted for the intraerythrocytic cycle, we would expect some overlap between the selected features obtained from these two cases. In fact, a significant overlap is observed between the selected proteins (around 50 %) from these two datasets; however, the match is not very high, possibly due to different tagging of the labels (Figure 3d). In the first data, the cells are labeled by the developmental stages (ring, trophozoite, gametocyte, and schizont), whereas in the second data, the cells are labeled by time (30hpi, 36hpi, 42hpi, and GCD). It is possible that there is variability in time among cells entering different stages.

### Protein-Protein interaction network analysis filterers out crucial protein in function

After selecting the features, we built protein-protein interaction networks based on these selected features in all three datasets. Considering the interactions and interdependencies between proteins, we created complex links between them by utilizing the STRING software. We chose the interaction type as text mining, experiments, databases, and co-expression in STRING^16^. For the first dataset, PPI network has 350 connected nodes, 1445 edges, average node degree of 8.26, avg. local clustering coefficient 0.386, PPI enrichment p-value: < 1.0e-16. For the second dataset, PPI network has 200 connected nodes, 287 edges, average node degree of 3.01, avg. local clustering coefficient 0.336, PPI enrichment p-value: < 1.0e-16. For the third dataset, PPI network has 239 connected nodes, 258 edges, average node degree of 2.16, avg. Local clustering coefficient 0.266, PPI enrichment p-value: < 1.0e-8. We calculated the degree and betweenness-centrality of each node and chose some proteins with high degree and betweenness centrality using Gephi^17^.Those nodes having degrees above 30 and betweenness centrality above 950 were slected and this set of proteins is denoted as A. Markov clustering(MCL) method is used to find different clusters in this network. We can see from the figure 4a that all the high degree and betweenness centrality proteins belong to one of these clusters. Same network analysis for selected features of the second dataset were performed and the nodes that have degrees above 15 and betweenness centrality above 500 were chosen. This set of proteins is denoted as B. For the third dataset, those nodes that have degrees above 15 and betweenness centrality above 500 were cosen. As the first two datasets are intraerothrocyte datasets, We chose common proteins from sets A and B. Gene ontology enrichment analysis with these selected features was performed using g:Profiler^18^ which is a popular toolset for converting gene identity mappings to orthologs and discovering biological categories enriched in gene lists. We observed that PF3D7_0810300, PF3D7_1105000, PF3D7_1246200 proteins(in asexual stages) are members of some enriched biological functions and we have marked all the enriched biological functions in figure 4b, 4c. A close observation of the gene ontology analysis suggests that the apical complex and extracellular proteins seem to be commonly enriched in both intraerythrocyte data sets pointing to plausible changes in the apical and extracellular components during the plasmodium life cycle stages. We followed the same method for the third dataset i.e., non-intraerothocyte dataset in the sexual phase.

**Figure 4.**
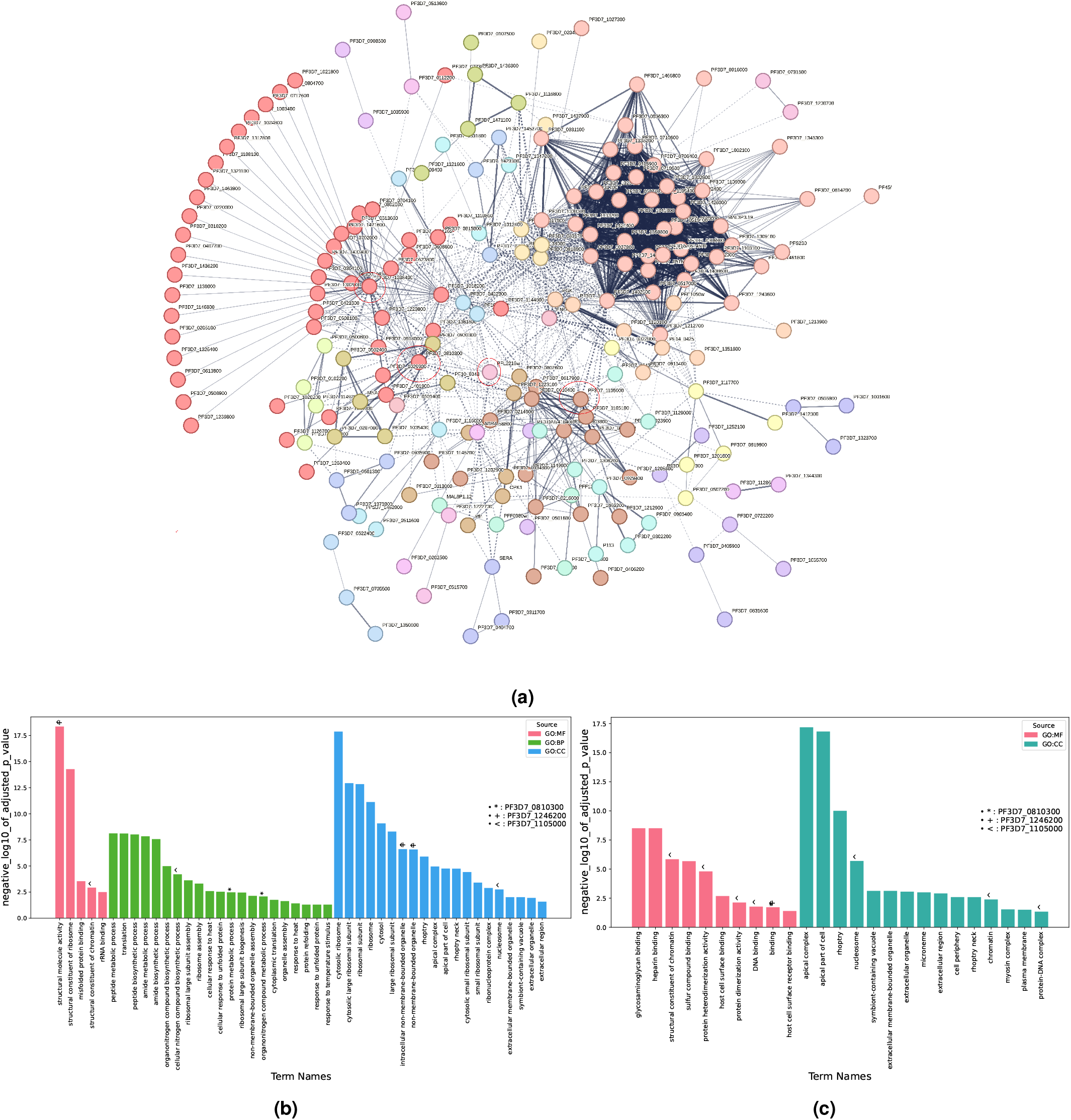
PPI and GO analysis identifies crucial proteins for drug targets. (a) Protein-protein interaction network constructed from STRING database for the selected features is described with clustering. Different colors represent different clusters. The selected four proteins for drug targets are highlighted in the figure by red circles. (b) Enriched biological functions of selected features with at least one connection in the PPI network for the first dataset. (c) Enriched biological functions of selected features with at least one connection in the PPI network for the second dataset.

### Function of crucial proteins

*Plasmodium falciparum* has both sexual and asexual blood stages. Most malaria drugs target asexual blood stages because replication of the parasite within red blood cells at this stage causes symptoms(fever and anemia) in humans^19^. Many antimalarial drugs(e.g., artemisinin and its derivative) target the proteins of this stage. On the other side, the sexual stage is responsible for transmitting gametocytes to mosquitoes^20^. Table 1 denotes the four crucial proteins for the asexual stages identified by our network analysis and the corresponding functions are described below

**Table 1.** Crucial proteins from the first and second dataset which have high degree and betweenness centrality.

| Protein name | Degree | Betweenness centrality |
| --- | --- | --- |
| PF3D7_1038400 | 55 | 5228.2 |
| PF3D7_0810300 | 38 | 2080.6 |
| PF3D7_1105000 | 31 | 1427 |
| PF3D7_1246200 | 22 | 953.4 |

Table 2 denotes the four crucial proteins for the sexual stages identified by our network analysis and the corresponding functions are described below

**Table 2.**
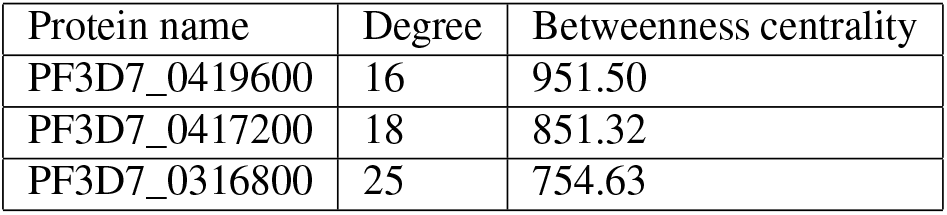
Crucial proteins from the third dataset which has high degree and betweenness centrality.

It is also desirable that if we can block gametocyte-specific proteins, we can control the spread of malaria. So, gametocyte-specific protein is also a good target to control malaria spread^20^. We have selected four proteins(check table 1) that are crucial for malaria in asexual stages, and we have analyzed their functions

- PF3D7_1038400: This is a Gametocyte-specific protein (Pf11-1) associated with the asexual stage of the Plasmodium life cycle. The P. falciparum gene Pf11-1 appears to code for a glutamic acid-rich protein that plays a similar role in the rupture of erythrocytic membranes, allowing gametes to escape, according to studies of mutant clones^21^. Therefore, sporogony and transmission would be prevented if this mechanism were disrupted. We can use this protein to find a suitable drug that can be used to prevent transmission.
- PF3D7_0810300: Protein phosphate catalyzes the removal of phosphate groups from proteins and regulates their activity. Therefore, Protein phosphatases play important roles in cellular processes. It helps in Oocyst development^22^. This protein does not have any strong binding sites.
- PF3D7_1105000: This protein is a Core component of nucleosome. Nucleosomes wrap and compact DNA into chromatin, limiting DNA accessibility to the cellular machinery, which requires DNA as a template. Thus, histones are essential for controlling transcription, repairing DNA, replicating DNA, and maintaining chromosomal stability. DNA accessibility is regulated via a complex set of post-translational modifications of histones, also called histone code, and nucleosome remodeling^23^.
- PF3D7_1246200: Plasmodium parasites have the actin-1 protein, which is a member of the actin family. Actin Plays an important role in the plasmodium life cycle. It has several functions:

Mobility and invasion: Highly conserved actin polymerizes to create filaments that create cytoplasmic cross-linked networks.^24^. Actin is necessary for the motility and invasive characteristics of Plasmodium. This protein is expressed in all the life stages of the parasite. Actin is involved in the motility of various life stages, including the invasive stages, such as sporozoites that need to move through tissues and invade host cells^25^.

Erythrocyte Invasion: Actin participates in the invasion of red blood cells in the blood stage of Plasmodium, which causes the symptoms of malaria. The parasite enters the host cell through the use of an actin-myosin motor^26^.

Cell Division: During asexual replication, actin is important in cell division of Plasmodium in the host’s red blood cells^25^.

Endocytosis and Vesicle Trafficking: Actin involves several intracellular processes, such as vesicle trafficking and endocytosis. These procedures are vital for the intake of nutrients and the production of proteins, both of which are necessary for the parasite’s survival and proliferation^27^.

We have selected three proteins that are crucial for malaria in sexual stages(check table 2) and we have analyzed their functions

- PF3D7_0419600: This is Ran-specific GTPase-activating protein 1. This putative RanGAP1’s particular role is likely to regulate Ran GTPase activity in a way similar to how it functions in other eukaryotes. This may affect various cellular functions, nuclear transport, and spindle formation during cell division. This protein does not have a strong binding site found^28^.
- PF3D7_0417200: Bifunctional dihydrofolate reductase-thymidylate synthase (DHFR-TS) is an enzyme that plays a crucial role in the folate biosynthesis pathway in many organisms, including Plasmodium parasites^29^. Here’s a brief overview of its functions:

Dihydrofolate Reductase (DHFR): The synthesis of purines, thymidylate, and certain amino acids depends on the conversion of dihydrofolate to tetrahydrofolate, which is carried out by this portion of the enzyme. The synthesis of amino acids and nucleic acids (DNA and RNA) requires tetrahydrofolate.^29^.

Thymidylate Synthase (TS): This portion of the enzyme synthesizes Thymidine, an essential component for DNA replication and repair. Thymidine is one of the four nucleotides that make up DNA.^29^.

The DHFR-TS enzyme is a well-known target of antimalarial drugs such as trimethoprim and pyrimethamine in Plasmodium and other species^30^. These drugs inhibit DHFR’s function, which throws off the parasite’s folate metabolism. This hinders the parasite’s capacity to multiply and thrive by making it unable to synthesise nucleic acids, such as DNA and RNA. Since nucleotide biosynthesis and folate metabolism are intertwined, the bifunctional nature of DHFR-TS in Plasmodium makes it a desirable target for antimalarial drug development. Notably, mutations in the DHFR-TS gene can result in resistance to antimalarial drugs, reflecting ongoing challenges in treating malaria.

- PF3D7_0316800: This is 40S ribosomal protein S15A. There is no strong binding site found in this protein.

### Drug Generation Using Deep Learning

In the final phase of our research, we turned our attention to drug discovery and generation. Predicting binding sites of crucial proteins marked the initiation of this stage, employing tools such as Cavity^31^. (figure 5a, 5b). Ligand binding sites offer crucial information to understand protein activities and structure-based drug design. We used CavityPlus software to make the property analysis and cavity detection process easier. The CAVITY programme is used by CavityPlus to identify putative binding sites in a given protein structure. In the asexual stage, we have identified four crucial proteins as described in the previous section. Out of those four, PF3D7_1038400 has a sequence length of 9563. Because of the long length, PF3D7_1038400 did not fit into the cavity^31^ for the binding site search. So, We have searched a 100 percent similar protein named A0A024W5F1 of sequence length 312 and searched its binding site, but no strong binding site was found in this protein. We did not get any strong binding sites for PF3D7_0810300 and PF3D7_1105000 proteins also. Only, PF3D7_1246200 protein displays strong binding sites. In the sexual stage, the PF3D7_0417200 protein was found to have five strong binding sites. We have used a non-autoregressive generative diffusion model, TargetDiff ( figure5c), to generate drug molecules targeting these strong binding sites^11^. As far as our knowledge, Targetdiff is the only target-based non-autoregressive diffusion model known today. Then we analyzed different properties of these drug molecules, such as physicochemical properties(molecular weight, volume, density, number of hydrogen atom acceptors(nHA), number of hydrogen atom donors(snHD), number of rotatable bonds in a molecule(nRot), Total Polar Surface Area(TPSA), Logarithm of Solubility (logS), Logarithm of the Partition Coefficient(logP), etc.), medicinal properties(Quantitative Estimate of Drug-likeness.(QED), Surface Area to Volume Ratio score(SAS), Lipinski rule, Pfizer rule, GSK rule, Golden Triangle rule, etc), ADMET properties, Toxicity using ADEMETlab 2.0^32^. We have chosen a few molecules that satisfy Lipinski, Pfizer, GSK, and Golden Triangle rules. We have used swissADME^33^ to find aleadlike molecule which satisfies drug-likeliness properties(Lipinski rule, Ghose filter, Veber rule, Egen rule, Muegge rule) and that has no problematic fragment in its structure^34–37^. Details of physicochemical properties and medicinal properties are given in the method. We have found these problematic fragments using PAINS (for pan assay interference compounds) and the Structural Alert Method^33^. Finally, we got few molecules that can be used as a drug against this protein, PF3D7_1246200 (figure (figure 6) and PF3D7_0417200 (check table 3). In order to verify the binding affinities of these drug molecules to the target,docking simulations were performed to obtain the binding energies and stable protein-ligand conformer (figure 6h) of each selected drug molecule with the respective target using Autodoc Vina.^38,39^. The binding energies of each drug molecule(ligand_i means i-th drug in the asexual stage in table 3 ) in the asexual stage with the respective target are given in figure 6g. The binding energy of the sable conformation shows high binding energy. This integrated approach provided a holistic understanding of potential drug candidates, considering their structural features and pharmacological properties, paving the way for further exploration and development of lead compounds for combating Plasmodium falciparum infections.

**Table 3.**
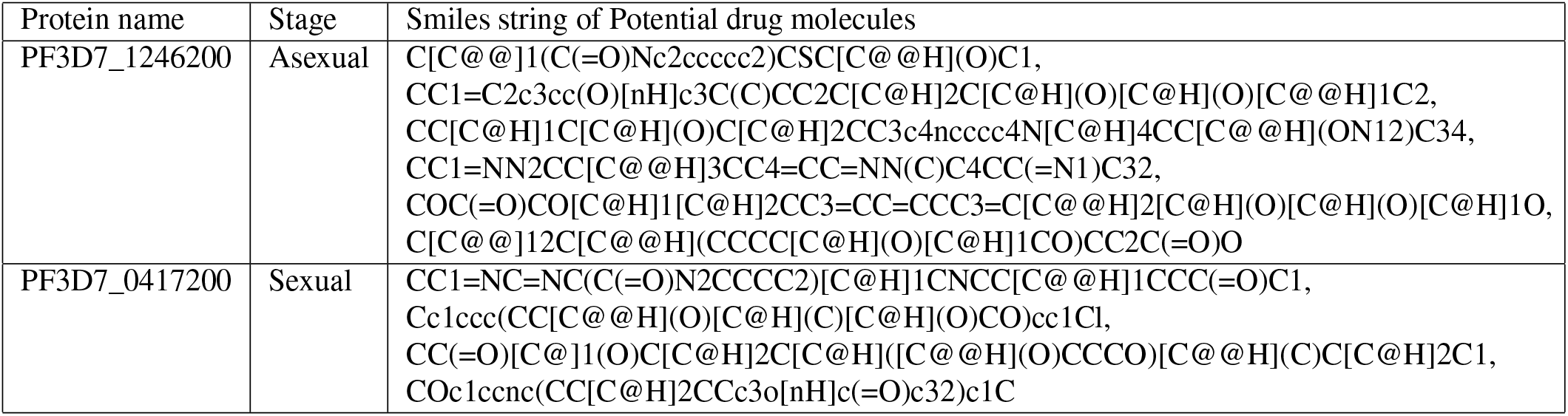
Selected drug molecules for PF3D7_1246200 and PF3D7_0417200.

**Figure 5.**
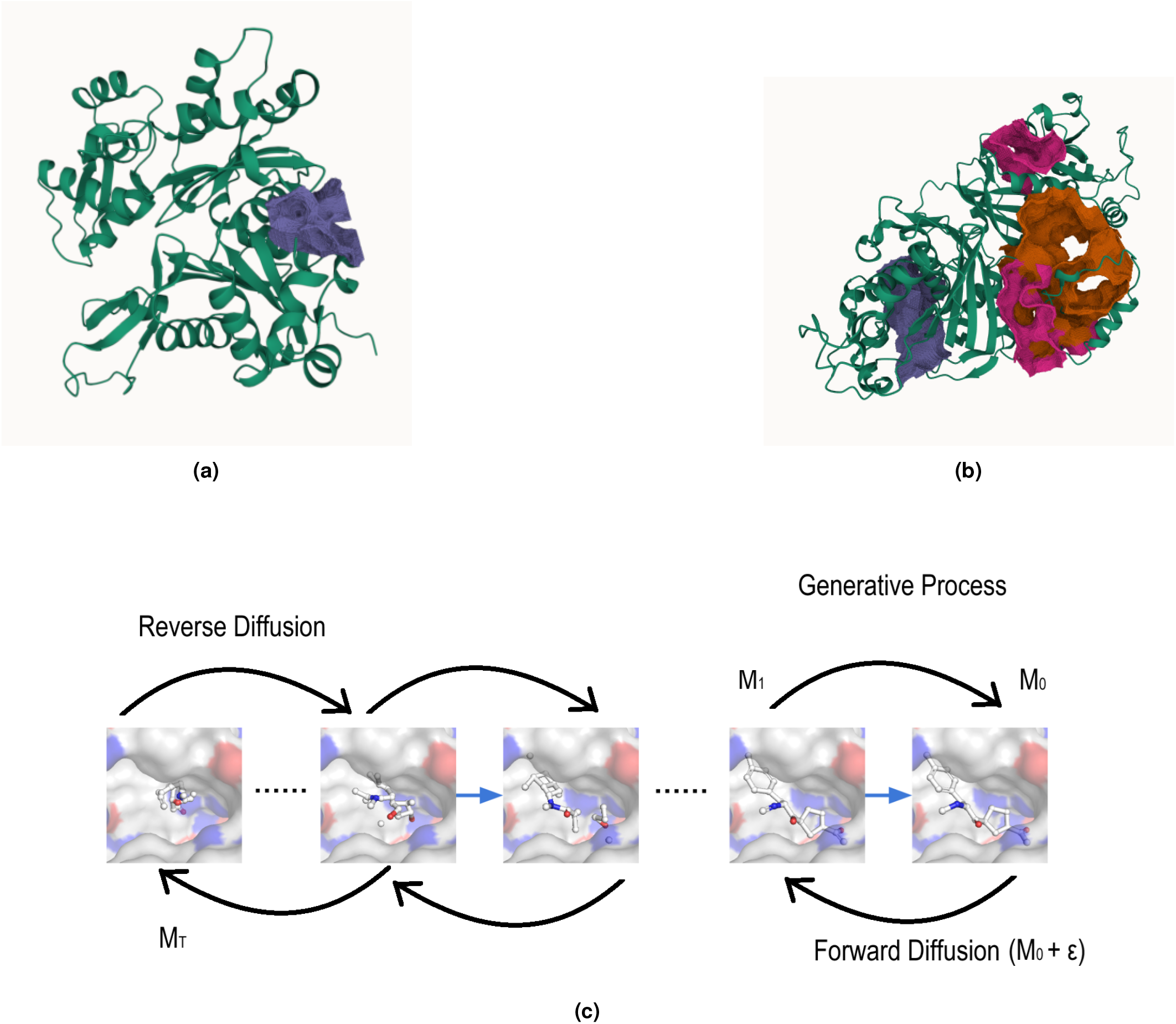
Binding site prediction and deep learning based diffusion model predicts potential drug molecules. (a) A 3D representation of one strong binding site of PF3D7_1246200 protein as identified by the CAVITY software (b) A 3D representation of one strong binding site of PF3D7_0417200 protein as identified by the CAVITY software. (c) The architecture of targetdiff diffusion model. *M*_0_ is the molecules in dataset. we are adding noise *ε* to *M*_0_ and getting a diffused Gaussian noise *M*_*T*_ . From *M*_*T*_ we are doing reverse diffusion and generating new samples.

**Figure 6.**
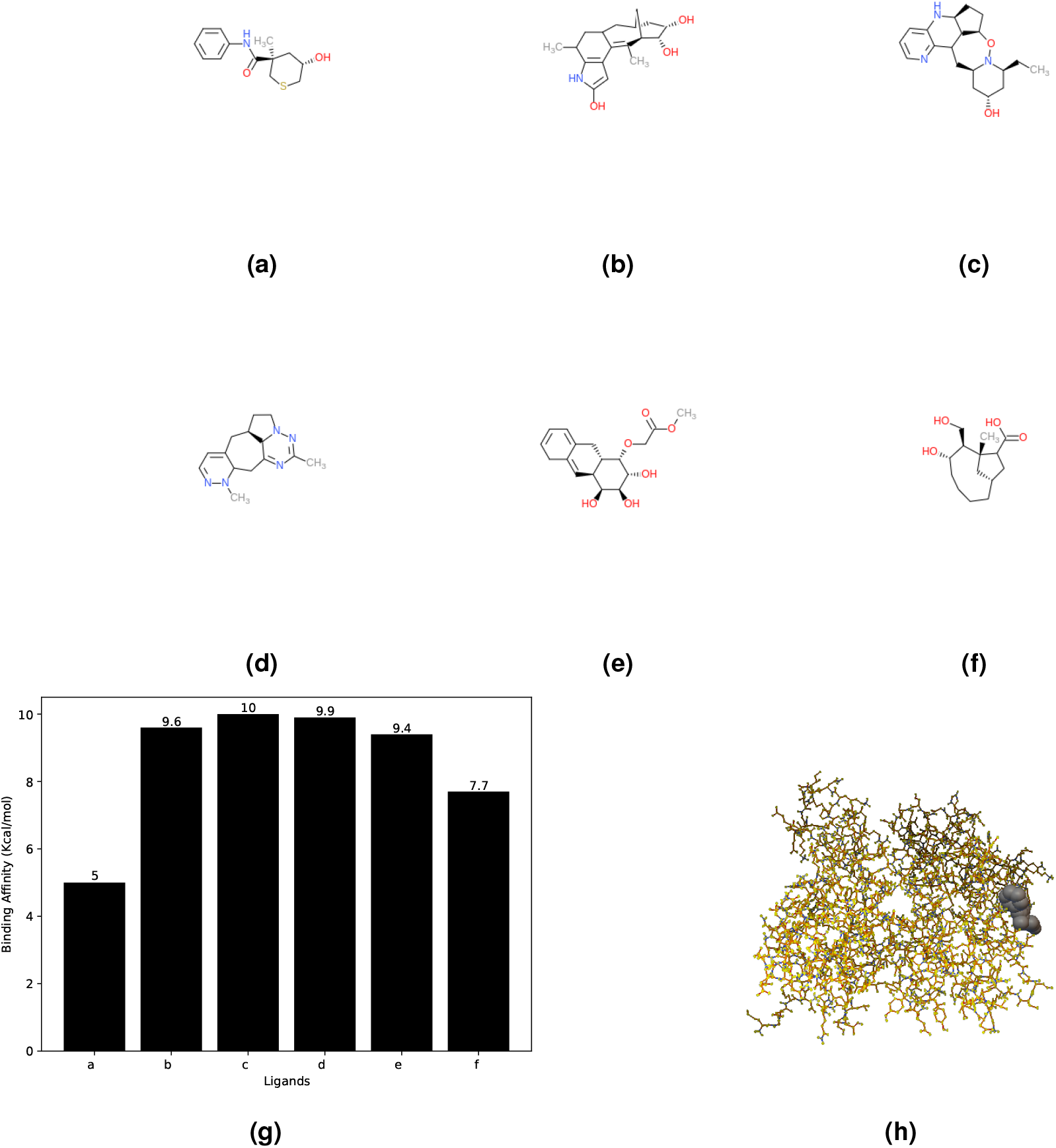
The targetdiff diffusion model generates a set of drug molecules for the PF3D7_1246200 protein. (a) The molecular formula for smile string representation of C[C@@]1(C(=O)Nc2ccccc2)CSC[C@@H](O)C1, (b) CC1=C2c3cc(O)[nH]c3C(C)CC2C[C@H]2C[C@H](O)[C@H](O)[C@@H]1C2, (c) CC[C@H]1C[C@H](O)C[C@H]2CC3c4ncccc4N[C@H]4CC[C@@H](ON12)C34, (d) CC1=NN2CC[C@@H]3CC4=CC=NN(C)C4CC(=N1)C32, (e) COC(=O)CO[C@H]1[C@H]2CC3=CC=CCC3=C[C@@H]2[C@H](O)[C@H](O)[C@H]1O, (f) C[C@@]12C[C@@H](CCCC[C@H](O)[C@H]1CO)CC2C(=O)O (g) represent comparisons of binding affinities between selected ligands in table 3 with its target protein for asexual stages. (h) represent stable protein-ligand stable binding complex for PF3D7_1246200 protein and 1st ligand(a).

## Discussion

Our study presents an important protocol for in silico drug design, aiming to identify crucial proteins that can be used as drug targets and generate drug molecules that can inhibit these proteins. We utilize three single-cell RNA-seq datasets covering distinct developmental stages of the parasite. Initially, we did data cleaning, normalization, and transformation. Several classification models e.g. Logistic Regression, Random Forest, and XGBoost—to predict blood cycle stages based on gene expression patterns were deployed in conjunction a mutual information-based feature reduction technique. We have selected these features in such a way that classification accuracy remains the same or increase compared to accuracy before feature selection. We used these selected features to construct protein-protein interaction networks to elucidate the molecular relationships governing the parasite’s developmental stages. The topology of network architecture, considering the degree and betweenness centrality of nodes, gave valuable insights into crucial proteins influencing interactions and information flow within the networks. We did biological functions enrichment analysis with the selected feature and found some crucial proteins that are significant for plasmodium survival. After finding strong binding sites cavity of those proteins, generative deep learning model ‘TargetDiff’ was employed to generate potential drug candidates^11^. Nowadays, Generative deep learning models are enhancing the drug design process effectively^40^. There are numerous deep learning-based based that have gone for clinical trials, such as Atazanavir(An FDA-approved antiviral drug used as a therapy for HIV discovered by Eargen’s MT-DTI^41^), Remdesivir ( An antiviral medicine under clinical trial discovered by Eargen’s MT-DTI^42^), Ruxolitinib (under clinical trials for COVID-19 discovered by Benevolent AI^43^), Baricitinib (got FDA permission for use with Remdesivir, resulting in a higher recovery rate for hospitalized COVID-19 patients discovered by Benevolent AI^44^), etc. Although the deep learning-based drug design paradigm is still in its infancy, such efforts would shed more light on the efficacy of the techniques and facilitate its growth towards maturity. We analyzed different properties of our generated drug molecules, such as physicochemical properties, medicinal properties and ADMET properties using ADEMETlab 2.0^32^ and have chosen a few molecules that satisfy Lipinski, Pfizer, GSK, and Golden Triangle rules. We have used swissADME^33^ to find a leadlike molecule that satisfies drug-likeliness properties and has no problematic structure fragment. Next, we did docking to find the binding energies and stable protein-ligand conformer of each selected drug molecule with the respective target using Autodoc Vina.^38,39^. Finally, the protocol provides us with a few potential drug molecules for the targets. This research is significant because it takes a comprehensive approach that combines target finding, network analysis, and drug development. Our study lays the groundwork for focused therapies against Plasmodium falciparum by identifying important genes, deciphering protein interactions, and finding viable drug candidates. The impact of the feature selection procedure on preserving or enhancing classification model accuracy highlights the method’s importance and confirms the chosen genes’ applicability in blood cycle stage prediction. Regarding the future direction, our discoveries set the foundation for new avenues in malaria research. There is potential for additional experimental validation and improvement with the discovered proteins and drug candidates. Furthermore, broadening the scope of the research to encompass larger datasets and integrating sophisticated machine-learning methodologies may improve our models’ accuracy and applicability. Validating the therapeutic potential of the selected drug candidates will need integrating experimental validations, such as in vitro assays and animal models. Furthermore, examining the wider relevance of our methodology to alternative parasitic illnesses may enhance the influence of our discoveries beyond *Plasmodium falciparum*. Our study provides a general framework that can be used for any single-cell data of any disease and hopefully would advance drug design research of *Plasmodium falciparum* .

## Method

We have used three datasets of single-cell transcriptomics. First data set is a rich atlas of short and long-read single-cell transcriptomes of over 37,000 Plasmodium falciparum cells across intraerythrocytic asexual and sexual development^12^. Second data set is Single-cell transcriptomes collected from early (ring) and late asexual blood stages, as well as stage I and stage IV gametocytes (GC)^13^. Third dataset is Single-Cell transcriptomics collected from the zygote and the ookinete stages^14^. First two datasets are intra-erothrocyte datasets, and third dataset is non intra-erothrocyte dataset. Each row of these datasets corresponds to a single cell and each column corresponds to a gene.

### Data prepossessing

The raw data collected from single-cell experiments require preprocessing to address various challenges and enhance the quality of downstream analyses. Data preprocessing in single-cell transcriptomics has a few steps that involve cleaning, normalizing, and transforming the raw data into a format suitable for meaningful biological interpretation^45^. We have done data cleaning for the three datasets to get a meaningful biological representation for our work. Variability in sequencing depth and library sizes amongst cells is a common difficulty that might lead to biases in the study. Total count normalization, which scales each cell’s expression values according to its total number of reads, is widely used to correct this. We have normalized our datasets. We have normalized each gene expression by dividing the sum of the counts of that gene counts over each single cell and again dividing that gene count by the length of that gene.

### Feature selection

Measurement errors, technical variations, and noise can all be present in single-cell data^46^. Irrelevant or noisy features in the analysis can lead to inaccuracies for further analysis. Feature selection helps in identifying and eliminating such noisy features. Selecting a subset of relevant features allows researchers to focus on biologically meaningful genes or proteins^47^. After getting the prepossessed datasets, we implemented classification models(Logistic regression, Random forest, XGboost) to these datasets and found accuracies of these models. To select important features from these datasets, We implemented mutual information-based feature reduction technique to select important features. This algorithm calculates mutual information between each column and the target column. There are other feature reduction techniques, but mutual information-based feature reduction is important because this algorithm quantifies the amount of information shared between different variables and is particularly useful for capturing non-linear relationships, which is essential in biological data where relationships may not be strictly linear. MI-based algorithms are less sensitive to noise than variance-based feature selection methods. Single-cell transcriptomics datasets are often heterogeneous, containing a variety of cell kinds and states. This heterogeneity can be accommodated by mutual information-based feature reduction, which finds relevant features for differentiating between different cell types or circumstances. The main target of this feature reduction technique is that we chose features so that after using these classification models, the accuracy remains the same or increases. We have also shown that if we randomly select the same amount of features from these datasets and apply classification algorithm, then accuracy will decrease. We can say that these selected features are crucial and contribute more to datasets. We have shown the accuracies of these classification models before feature selection, after feature selection and randomly selected features in this bar diagram.

### Construction and analysis of protein-protein interaction network

Protein-protein interaction network is the interaction network between proteins where proteins are the nodes and edges are defined as the interaction between proteins. Since protein-protein interactions (PPIs) are necessary for almost all cellular processes, an understanding of PPIs is essential to understanding cell physiology in both normal and pathological situations^48^. According to Graph theory, PPI network’s topological structure gives relevant connections and information regarding the associated biological functions^49^. In network theory, degree(D), Connectivity degree (k), betweenness centrality (BC), closeness centrality (CC), eigenvector centrality (EC), and eccentricity are the fundamental measures^4^. Degree of a node denotes the number of edges connected with that node. Betweenness centrality of a given node is given by

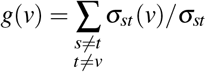

here, *σ*_*st*_ is the total number of shortest paths from node from s to t and *σ*_*st*_(*v*) is the number of those paths that pass through v and v is not the endpoint. Betweenness centrality measures the extent to which a node lies on the shortest paths between other nodes, indicating its significant influence over information flow. Nodes with high BC, called bottlenecks, tend to indicate essential genes because they can be compared to heavily trafficked intersections on major highways or bridges. In a PPI network, nodes with large degrees, referred to as hub proteins, may correspond to the genes that cause the disease. The hubs and bottlenecks at the heart of the PPI network were the primary focus of this investigation. In our work, after getting selected features, we constructed protein-protein interaction network with these selected features from three datasets. We have used string software to construct these networks^16^. Then, from these networks, the degree and betweenness centrality of all the nodes of the networks were obtained, and a few proteins with high degree and betweenness centrality are chosen as targets. These proteins have more control in this network. We performed gene ontology enrichment analysis with these selected features using g:Profiler^18^, a popular toolset for converting gene identity mappings to orthologs and discovering biological categories enriched in gene lists. It is found that all the crucial proteins are members of some enriched biological functions. A close observation suggests that the apical complex and extracellular proteins are commonly found in both the datasets suggesting the critical roles of these complements during the intraerythrocyte developmental phases.

### Binding site prediction and drug molecule generation

After doing PPI analysis, we selected a few proteins that are hubs. Then, we found the function of each protein and its role in plasmodium falciparum survival. Among these proteins, we select a few crucial ones that are important for plasmodium falciparum survival. We chose those proteins as a drug target for malaria. We have found the binding site of these proteins. We have used Cavity^31^ to find the strong binding sites of these proteins and after getting the binding sites, we have used a generative deep learning model to generate suitable drugs that can bind with these binding sites (we have shown all selected drug molecules for asexual stage in figure 6a-6f). We have used targetdiff, a target-based generative diffusion model, to generate drug molecules^11^. Many deep learning models can be used to generate drugs, but this is a target-based non-autoregressive diffusion model that can generate bonds and atoms of the molecules simultaneously^50^. A diffusion process is a continuous-time stochastic process that involves a series of diffusion steps to convert a given data distribution into a more simple distribution. In every step, the data is progressively transformed into a target distribution by adding noise to it. In reverse diffusion, we predict the added noise in each step to get the previous sample. Reverse diffusion allows to generation of new samples from the distribution^50^ (Figure 5c). This model has performed better than other models in target-based drug discovery. We have generated 100 drug molecules for each strong binding site.

### Analysis of drug molecules to find lead compounds

A well-balanced combination of safety, pharmacokinetics, and biochemical behavior is what makes a drug successful. A drug candidate’s ability to achieve a desirable absorption, distribution, metabolism, excretion, and toxicity (ADMET) profile is as important to its success as having high potency and selectivity. More precisely, the perfect drugs should be ingested into the body, distributed to different tissues and organs reasonably, digested so as not to erase their action, and removed suitably instantly, and their non-toxicity should be established. These concerns, which span the entire process from administration to elimination, appear separate but are intimately related. We analyzed different properties of these drug molecules, such as physicochemical properties(molecular weight, volume, density, number of hydrogen atom acceptors(nHA), number of hydrogen atom donors(snHD), number of rotatable bonds in a molecule(nRot), Total Polar Surface Area(TPSA), Logarithm of Solubility (logS), Logarithm of the Partition Coefficient(logP), etc.), medicinal properties(Quantitative Estimate of Drug-likeness(QED), Surface Area to Volume Ratio score(SAS), Lipinski rule, Pfizer rule, GSK rule, Golden Triangle rule, etc), ADMET properties using ADEMETlab 2.0^32^. Drug distribution, metabolism, excretion, and absorption are all impacted by molecular weight (ADME). High molecular weight may impact bioavailability and penetration of biological barriers^51^. Molecular packing, which affects drug stability and solubility, is impacted by volume. It can affect the transport of the drug via biological membranes^35,52^. The molecule’s capacity to create hydrogen bonds, which is essential for interactions with biological targets and solubility in water, is influenced by the number of hydrogen atom donors (nHD) and acceptors (nHA) in the molecule^35,53^. The number of rotatable bonds (nRot) affects the flexibility of the molecule and how it interacts with target proteins. Drug metabolism may be impacted by an excess of rotatable bonds^35^. Total Polar Surface Area (TPSA) offers information about the polarity of a molecule, which influences how well it interacts with proteins and can pass through biological membranes^33,35^. The likelihood of a drug dissolving in water is indicated by the logarithm of solubility, or logS. Bioavailability depends on solubility because poorly soluble medicines may have poor absorption^53,54^. The partition coefficient, or logP, measures a molecule’s lipophilicity and how well it can pass across cell membranes. It is necessary to predict the distribution of drugs in tissues^55^. The Quantitative Estimate of Drug-likeness, or QED, predicts how closely a molecule’s characteristics match those of successfully used drug molecules^56^. The Synthetic accessibility score(SAS) is a metric that quantifies how easily a given molecular structure can be synthesized^33^. Lipinski’s rule states that a drug-like molecule should have no more than five hydrogen bond donors (the total number of nitrogen–hydrogen and oxygen–hydrogen bonds), no more than ten hydrogen bond acceptors (all nitrogen or oxygen atoms), molecular mass should be less than 500 daltons, log P should be less than five^53^. Pfizer Rule states that a drug-like molecule should have logP greater than 3 and TPSA less than 75^57^. GSK Rule states that a drug-like molecule should have molecular weight ≤ 400 and logP ≤ 4^58^. Golden Triangle Rule states that a drug-like molecule should have 200 ≤ molecular weight ≤ 50; -2 ≤ logD ≤ 5^59^. After that, we used swissADME^33^ to find a leadlike molecule that satisfies drug-likeliness properties(Lipinski rule, Ghose filter, Veber rule, Egen rule, Muegge rule) and that has no problematic fragment in its structure. We have found these problematic fragments using PAINS (for pan assay interference compounds) and the Structural Alert Method^33^. Ghose Filter states that a leadlike molecule should have a molecular weight between 160 and 480 daltons, LogP between -0.4 and +5.6, Number of atoms between 20 and 70, molar refractivity between 40 and 130^34^. Veber Rule states that a leadlike molecule should have No more than 10 rotatable bonds, No more than 140 Å^2^ polar surface area^35^. Egan Rule suggests that a compound is likely to be orally active if it has logP between 1 and 3, a molecular weight between 150 and 500 daltons^36^. Muegge Rule suggests that a leadlike molecule should have No more than 5 hydrogen bond donors, No more than 10 hydrogen bond acceptors, No more than 10 rotatable bonds, A polar surface area of less than 140-angstrom square.^37^. We have chosen a few drug molecules that satisfy lead-likeliness and drug-likeliness properties and have no problematic structure fragments. After generating drugs we filter these drug molecules with the ADMET, druglikeliness, and leadlikeliness properties. Finally, after filtration we selected some drug molecules for each protein (table 3). Next, we performed docking to find the binding energies and stable protein-ligand conformer of each selected drug molecule with the respective target using Autodoc Vina.^38,39^.

## Acknowledgements

Authors thank Department of Biotechnology (No. BT/RLF/Re-entry/32/2017), Government of India for funding this project.

## Author contributions statement

Conceptualization: S.C. B.G.; methodology: S.C.; formal analysis and investigation: S.C. Writing—original draft preparation: S.C.; writing—review and editing: S.C. B.G.; funding acquisition: B.G.; supervision: B.G.

## Additional information

**Supplementary Information** : All the codes and related supplementary material available at (https://github.com/soham2-4/Malaria-Drug-Targets-finding-from-Single-Cell-Transcriptomics-and-genertaing-potential-drug-molecule).

